# TGF-β and BMP subfamily pathways in zebrafish spermatogonial niche: A83-01 and DMH1 inhibitor effects in spermatogonial self-renewal and differentiation

**DOI:** 10.1101/2022.10.21.513182

**Authors:** Daniel Fernandes da Costa, Juliana Morena Bonita Ricci, Maira da Silva Rodrigues, Marcos Antônio de Oliveira, Lucas Benites Doretto, Rafael Henrique Nóbrega

**Affiliations:** Reproductive and Molecular Biology Group, Department of Structural and Functional Biology, Institute of Biosciences, São Paulo State University (UNESP), 18618-970, Botucatu, São Paulo, Brazil; South Bohemian Research Center of Aquaculture and Biodiversity of Hydrocenoses, Research Institute of Fish Culture and Hydrobiology, Faculty of Fisheries and Protection of Waters, University of South Bohemia in Ceske Budejovice, 389 25 Vodňany, Czech Republic

**Keywords:** TGF-β, BMP, A83-01, DMH1, spermatogonia, self-renewal, differentiation

## Abstract

This study unravels the roles of TGF-β (Transforming growth factor-β) superfamily signaling pathway in the zebrafish spermatogonial activity (self-renewal *vs*. differentiation) by combining *ex vivo* and specific pathway inhibitors approaches. The TGF-β superfamily signaling pathway is subdivided into TGF-β and Bone morphogenetic proteins (BMP) subfamilies, and is ubiquitous among metazoans, regulating several biological processes, including spermatogenesis. In this study, we evaluated the function of the TGF-β and BMP subfamily pathways in the zebrafish spermatogonial niche using A83-01 and DMH1 inhibitors, respectively. Our results showed that A83-01 potentiated the follicle-stimulating hormone (Fsh) effects on zebrafish spermatogenesis, reducing type A undifferentiated spermatogonia and increasing differentiated spermatogonia (type Adiff and type B spermatogonia) after 7 days of culture. In agreement with histomorphometrical data, the mRNA levels of *dazl* (marker of spermatogonial differentiation) and pro-differentiation growth factors, such as *igf3* and *insl3,* were significantly augmented following A83-01. For the BMP signaling pathway, exposure to DMH1 inhibitor showed opposite effects as compared to TGF-β superfamily signaling pathway inhibitor. Histomorphometrical analysis demonstrated an accumulation of type A undifferentiated spermatogonia, while the frequency of differentiated spermatogonia was significantly reduced following co-treatment of DMH1 with Fsh after 7 days of culture. To support this data, expression analysis revealed that BMP signaling pathway inhibitor also decreased the testicular mRNA levels of *dazl, igf3* and *insl3* when compared to control incubation (Fsh). In conclusion, our study demonstrated that TGF-β and BMP subfamily pathways exert a role in zebrafish spermatogonial niche with antagonistic functions for the spermatogonia fate. The TGF-β subfamily pathway is involved with spermatogonial self-renewal and inhibition of differentiation, whereas the BMP subfamily pathway promotes spermatogonial differentiation. These findings are not only relevant to understanding stem cell biology, but may also be useful in several in vitro assays, promoting control of self-renewal and differentiation by potentially directing these processes.

## 1. Introduction

Transforming growth factor β (TGF-β) superfamily signaling pathways are essential regulators of cellular processes including proliferation, differentiation, survival, as well as key players for different biological processes such as embryonic development, angiogenesis, regeneration and gametogenesis [1–6]. The TGF-β superfamily signaling pathway is activated by TGF-β superfamily members, which comprise dozens of autocrine and paracrine regulatory ligands. Some of these ligands are known by multiple names according to the context of their discovery [1,4,7,8]. They function as dimers and each subunit contains seven conserved cysteine residues that form three intramolecular disulfide bonds and one inter-subunit disulfide bond [1,7].

The TGF-β superfamily signaling pathway is evolutionarily conserved in metazoans [9]. The principal TGF-β superfamily members are TGF-βs 1-3 (encoded by different genes); nodal; activins (formed by activin β subunit dimers, also known as inhibin β subunits); inhibins (formed by inhibin α and inhibin β subunits); bone morphogenetic proteins (BMPs); anti-Mullerian hormone (AMH); growth and differentiation factors (GDFs); and glial cell line-derived neurotrophic factor (GDNF) [1,7,10–12]. Several proteins of TGF-β superfamily members have been found in different species, such as *Xenopus laevis* [13], *Caenorhabditis elegans* [14], *Drosophila melanogaster* [15], *Danio rerio* [16,17], among others.

In general, TGF-β superfamily proteins are present in the extracellular matrix and bind to complexes of serine/tyrosine kinase type I and type II membrane receptors [3]. Type II receptors have an extracellular domain for specific association with ligands of the TGF-β superfamily, comprising the following receptors, TGF-βRII for TGF-βs; ActRIIA/ActRIIB for nodal, activin, and inhibin; BMPR-II and ActIIA/ActRIIB for BMPs; and AMRH-II for AMH [3,8,11,12]. Type I receptors act as downstream components of type II receptors in the signaling pathway and are denominated as ALKs (activin receptor-like kinases). ALKs 4, and 7 form a complex activated by nodal and activin, ALK 5 by TGF-βs, and ALK2, 3, and 6 by BMP and AMH [12,18]. ALK1 has restrictive expression (e.g., in endothelial cells) and is a promiscuous receptor that binds preferentially to BMPs as well as TGF-βs [19]. Following the formation of the ligand/receptor II and I complex, a cascade of phosphorylations, initiated with the type II receptor phosphorylating the type I receptor, activates cytoplasmatic mediator proteins called Smads [18,20]. Smads are divided into three groups according to their functions. R-Smads (Smads1, 2, 3, 5, and 8) are associated with type I receptors and mediate membrane signals to the nucleus. Co-smad (Smad4), which binds R-Smads in the cytoplasm, transporting them to the nucleus. Finally, I-Smads (Smads6 and 7) are antagonists of R-Smads and exhibit inhibitory activity under specific stimuli [1,7,12].

The TGF-β superfamily pathways are divided in two large subfamilies according to the ligands, type I receptors, and Smad mediators. The TGF-β subfamily uses type I receptors (ALK4, ALK5, and ALK7), is activated by the ligands TGF-βs, nodal, activin and inhibin and its signaling is mediated by R-Smad 2 and 3; and the BMP subfamily uses type I receptors (ALK2, ALK3, and ALK6), is activated by the ligands BMPs and AMH and its signaling is mediated by R-Smad 1, 5, and 8 [21]. In both pathways, Smad4 carries the phosphorylated R-Smads (1, 2, 3, 5, and 8) from the cytoplasm to the nucleus; this complex carried to the genomic DNA acts in the regulation of gene expression, in association with coactivators, corepressors, and other transcriptional regulators [7,11,20,22].

Among different species of fish, it has been shown that several reproductive processes, such as sex determination [23], sex differentiation [24], gametogenesis [23], fertilization and sexual behavior [25], are dependent of TGF-β superfamily signaling pathway. On the other hand, the role of TGF-β superfamily signaling pathway on fish spermatogenesis remains poorly investigated. A valuable approach to understand TGF-β superfamily signaling pathway is the use of specific inhibitors of signaling pathways. Several inhibitors of TGF-β type I receptors and BMP type I receptors have been developed (e.g., SB-431542, A83-01, dorsomorphin, and DMH1) [21,26–29]. Among these inhibitors, A83-01 blocked type I receptors in mink lung epithelial (Mv1Lu) cells [26], human mammary epithelial cells (HMLE) [30] and the human breast cancer cell line SKBR3 [31]. DMH1 blocked BMP type I receptors in non-small cell lung cancer (NSCLC) [29], induced human pluripotent stem cells (hiPSCs) [28] and zebrafish embryos [27].

In the present study, we investigated the two large subfamilies of TGF-β superfamily pathways (TGF-β and BMP subfamily) in zebrafish testes, focusing on the spermatogonial phase. For that, we pharmacologically inhibited the signaling pathway of TGF-β and BMP subfamily using A83-01 and DMH1 inhibitors, respectively, using zebrafish testis tissue cultures under basal conditions and following stimulation with Fsh (Follicle stimulating hormone), which is the main endocrine player regulating spermatogonial phase in zebrafish [32]. In order to clarify the possible roles of these signaling pathways in the regulation of the spermatogenic process (self-renewal or differentiation), we performed quantitative analyses of germ cells (type A undifferentiated spermatogonia, Aund; type A differentiated spermatogonia, Adiff, and type B spermatogonia), evaluation of expression of germ stem cell *(pou5f3, nanog)* and differentiation markers *(dazl),* growth factors *(amh, igf3, insl3),* and mediators of TGF-β and BMP subfamily pathways *(smad2, smad3a, smad3b, smad1, smad8*). The unprecedented comparison between the A83-01 and DMH1 inhibitors used in this study provided valuable information regarding the TGF-β and BMP subfamily signaling pathways and their roles in the spermatogenic process in zebrafish. Likewise, the functional evaluation between the Fsh and the TGF-β superfamily signaling pathways added important information regarding these signaling pathways and their influence on the spermatogonial niche of teleosts.

## 2. Material and methods

### 2.1 Animals

Adult male zebrafish (outbred) were bred and raised in the aquarium facility at the Department of Structural and Functional Biology, Institute of Biosciences, São Paulo State University (Botucatu, Brazil). Fish were kept in tanks of 6-L in a water recirculation system under constant temperature (28 °C) and controlled photoperiod (14 h light and 10 h dark). Salinity, pH, dissolved oxygen and ammonia were monitored daily. Fish were fed twice a day with the commercial food (Zeigler®). Handling and experimentation were consistent with Brazilian legislation regulated by the National Council for the Control of Animal Experiments (CONCEA) and Ethical Principles in Animal Research (Protocol n. 666-CEUA).

### 2.2 Testis tissue culture

Testes were dissected and cultured using an *ex vivo* organ culture system as previously described [33]. For short-term incubations (18 h for gene expression analysis), testes were submerged in a culture medium, whereas for long-term exposures (7 days for gene expression analysis and morphology), testes were placed on a nitrocellulose membrane on top of a cylinder of agarose and cultivated with 1mL of medium in 24-well flat-bottom plates, as previously described [33]. In all treatments with A83-01 and DMH1, DMSO (0,001% v/v) was administered to the respective controls.

### 2.3 Determination of the A83-01 and DMH1 inhibitor concentrations

To block TGF-β type I receptors, 1 μM A83-01 (Sigma-Aldrich, USA) was used, as shown in a previous study [26]. To determine the DMH1 (Sigma-Aldrich, USA) concentration required to block BMP type I receptors, a concentration test was performed. One testis was cultivated in Lebovitz medium (L-15) (Sigma-Aldrich, St. Louis, MO, USA), whereas its contralateral testis was cultivated in L-15 containing three different concentrations of DMH1 (1, 5, and 10 μM) for 18 h. For this experiment, eight adult males were used per concentration. Testes were incubated in 96-well plates containing 200 μL of solution in each well at 28°C. Following incubation, the testes were stored at −80°C to quantify *id1* (Inhibitor of DNA binding 1) gene expression; it is used to validate DMH1 inhibitor.

### 2.4 Short-Term (18 h) Incubation

After dissection, one testis was cultivated in L-15 (Sigma-Aldrich), whereas the contralateral testis was cultivated in L-15 containing A83-01 (1 μM) or DMH1 (1 μM). For this experiment, eight adult males were used for each inhibitor, in a total of 16 animals. To verify the influence of A83-01 and DMH1 inhibitors on the effects induced by Fsh, one testis was cultivated in L-15 with 100 ng/mL of recombinant zebrafish (rzf) Fsh according to previous studies [32] whereas the contralateral testis was incubated with rzf Fsh (100 ng/mL) in the presence of A83-01 (1 μM) or DMH1 (1 μM). For this experiment, eight adult males were used for each treatment, in a total of 16 animals. To verify the expression of *smads* and *id1* following rzf Fsh stimulation in the absence of inhibitors, adult testes (n = 8) were incubated with 100 ng/mL rzf Fsh and their contralateral ones in L-15 medium. All testes were incubated in 96-well plates containing 200 μL of the solution in each well at 28°C. Following incubation, the testes were stored at −80°C for gene expression analysis.

### 2.5 Long-Term (7 Days) Incubation

To study the effects of A83-01 and DMH1 inhibitors on zebrafish spermatogenesis in the presence or absence of Fsh stimulation. Two sets of experiments using long-term incubations were carried out to evaluate gene expression (n = 32 males) and for histomorphometrical analysis (n = 32 males). After dissecting the testes (paired structure), each testis (left and right) was placed on a nitrocellulose membrane measuring 0.25 cm^2^ (25 μm of thickness and 0.22 μm of porosity) on top of a cylinder of agarose (1.5% w/v, Ringer’s solution—pH 7.4) into a 24-well plate containing 1 mL of culture medium. In this system, one testis (left) was incubated in the presence of A83-01 (1 μM,n = 16) or DMH1 (1 μM, n = 16) and its contralateral testis (right) in basal culture medium (L-15). Similarly, one testis (left) was incubated in the presence of A83-01 (1 μM) and 100 ng/mL rzf Fsh (n = 6) or DMH1 (1 μM) and 100 ng/mL rzf Fsh (n = 16) and its contralateral testis (right) in a basal culture medium (L-15) with rzf Fsh (100 ng/mL). All media were changed every 3 days of culture. After 7 days, testes were collected for histomorphometric and gene expression analysis. Total RNA was extracted from testis tissue and the relative mRNA levels of *pou5f3, nanog*, *dazl*, *amh*, *igf3*, *insl3*, *smad1*, *smad2*, *smad3a*, *smad3b*, *smad8*, and *id1* were evaluated as described below (qPCR).

### 2.6 Gene Expression Analysis by Real-Time, Quantitative PCR (qPCR)

Total RNA from testes (short-term and long-term incubations) was extracted using a commercial RNAqueous®-Micro kit (Ambion, Austin, TX, USA), according to the manufacturer’s instructions. cDNA was synthesized as previously described [34]. qPCR reactions were conducted using 10 μL 2x SYBR-Green Universal Master Mix, 2 μL of forward primer (9 mM), 2 μL of reverse primer (9 mM), 1 μL of DEPC water, and 5 μL of cDNA. The relative mRNA levels of *pou5f3* (POU domain, class 5, transcription factor 3), *nanog* (nanog homeobox), *dazl* (deleted in azoospermia-like), *amh* (anti-Mullerian hormone), *igf3, insl3, smad1, smad2, smad3a, smad3b, smad8,* and *id1* (inhibitor of differentiation-1) were evaluated in the different experiments. The mRNA levels of the targets (Cts) were normalized to the reference gene β-actin and expressed as relative values of the control group (as fold induction), using the 2^-(ΔΔCT)^ method. Primers (Table 01) were designed based on the zebrafish sequences available in GenBank (NCBI, https://www.ncbi.nlm.nih.gov/genbank/).

### 2.7 Histomorphometrical analysis

Zebrafish testis tissue was fixed in Karnovsky (glutaraldehyde 4% paraformaldehyde 8% in phosphate buffer pH 7.2) overnight, dehydrated, embedded in Technovit (7100-Heraeus Kulzer, Wehrheim, Germany), sectioned at 3μm thickness (non-serial sections), and stained with 0.1% toluidine blue to quantify the different germ cell types at 100x objective using a high-resolution light microscope (Leica DM6000 BD, Leica Microsystems, Wetzlar, Germany). For morphometric analysis, eight adult males were used for each treatment (A83-01 or DMH1) and co-treatment (A83-01 + 100 ng/mL rzf Fsh or DMH1 + 100ng/mL rzf Fsh). One hundred histological fields from each animal (n = 32) were randomly selected to count the frequency of germ cell cysts (type A undifferentiated spermatogonia, type A differentiated spermatogonia and type B spermatogonia). Germ cells were identified according to morphological criteria established for zebrafish germ cells, such as size of the nucleus, number of nucleoli, chromatin condensation, and number of germ cells per cyst [35–37]. To this end, a Leica DMI6000 microscope was used.

### 2.8 Statistical Analysis

All data were subjected to normality Shapiro-Wilk test, which was followed by the Bartlett homogeneity variance test. Data are presented as mean ± SEM (standard error of the mean) and significant differences were identified using Student’s t-test (paired) (p < 0.05). Comparisons of more than two groups were performed using one-way ANOVA followed by the Student–Newman–Keuls test (p <0.05). All data analyses were performed using GraphPad Prism software (version 4.0; GraphPad Software, Inc., San Diego, CA, USA, http://www.graphpad.com).

## 3. Results

### 3.1 Determination of DMH1 inhibitor concentrations

To determine which concentration of DMH1 can inhibit the BMP subfamily pathway, we measured the relative mRNA levels of *id1* in zebrafish testes incubated with different concentrations of DMH1. *id1*gene is a direct and early response BMP subfamily target that encodes a protein that negatively regulates the basic helix-loop-helix proteins (Hollnagel et al., 1999; Katagiri et al., 2002 [38,39]. The *id1* promoter has specific sites (BMP-responsive element, BRE) to bind phosphorylated Smad1, 5, and 8 in association with Smad4; upregulating *id1* expression [40–42]. Thus, we performed a test with three different concentrations of DMH1 (1, 5, and 10 μM), showing that only the smaller concentration (1 μM) inhibited *id1* gene expression (Figure 1).

**Figure 1.**
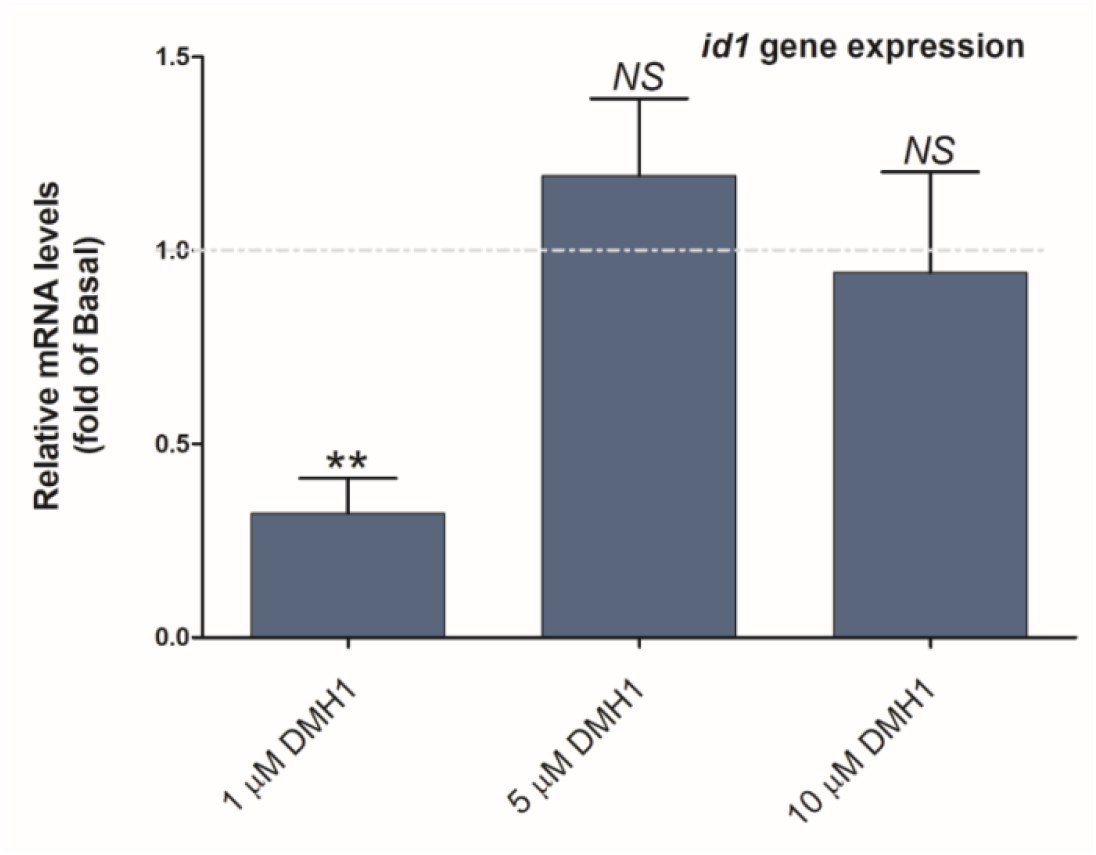
*id1* gene response to different concentrations of DMH1. Relative expression of *id1* gene after 18 h of exposure to different concentrations of DMH1 (1 μM, 5 μM, 10 μM) in zebrafish testis tissue culture. (**) represents significant difference of (p < 0,01). (NS) no significant difference. Data are presented as the mean ± standard error (n = 8).

### 3.2 Effects of the A83-01 and DMH1 inhibitors on gene expression of testis tissue cultured under basal condition

We evaluated the effects of the pharmacological inhibitors A83-01 and DMH1 in zebrafish testis tissue cultured under basal conditions (L-15) for 18 h and 7 days. After 18 h of cultivation with inhibitors, changes in expression were observed (Figures 2, 4). Treatment with DMH1 for 18 h downregulated all genes analyzed (Figures 2, 4), showing a tendency to repress the spermatogenic process. Treatment with A83-01 for 18 h elevated the relative expression of spermatogonial stem cell genes (Aund*/Aund), *pou5f3,* and *nanog*, in relation to basal culture conditions (Figures 2, 4). Conversely, after 18 h of A83-01 exposure, the levels of *igf3* and *smad3b* primary transcripts decreased and we found an inversely proportional ratio between *amh* and *igf3* expression (Figures 2, 4). However, the increase in *amh* was not significant (Figure 2).

**Figure 2.**
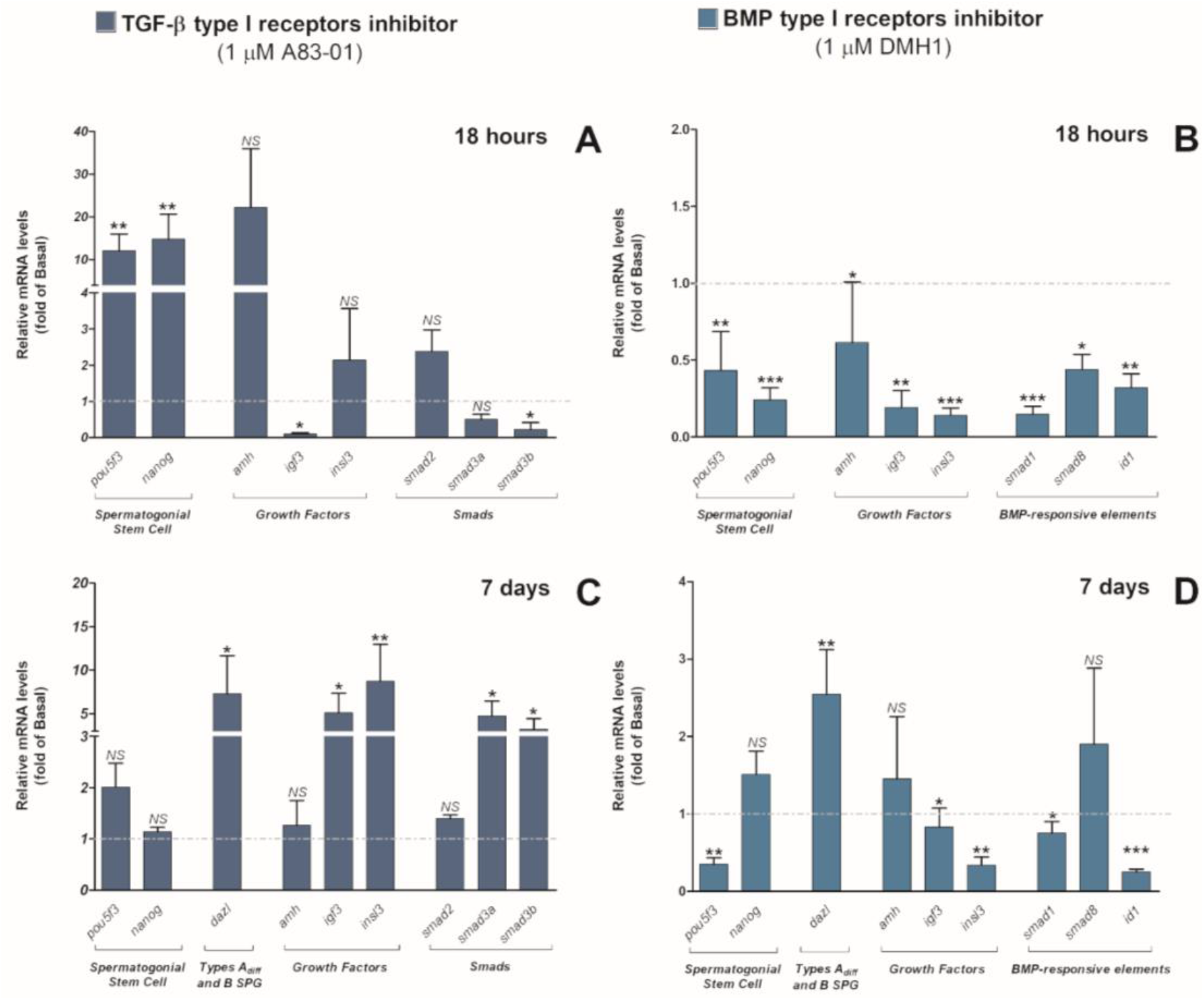
TGF-β and BMP subfamily pathway inhibition effects on gene expression in zebrafish testis tissue cultured under basal conditions. (A-D) Relative expression of spermatogonial stem cell (*pou5f3, nanog*), spermatogonial differentiation (*dazl*), growth factors (*amh, igf3, insl3*) markers, and intracellular messengers of the TGF-β superfamily pathway *(smad1, smad2, smad3a, smad3b, smad8, id1*) in testis tissue cultures treated with the A83-01 inhibitor (1 μM) for 18 h (A) and 7 days (C). DMH1 inhibitor (1 μM) after 18 h (B) and 7 d (D) of culture. Ct values were normalized to the reference gene *(β-actin)* and expressed relative to the basal group (L-15 only). (*), (**), and (***) represent significant differences of (p < 0,05), (p < 0,01), (p < 0,001), respectively. (NS) no significant difference. Data are presented as the mean ± standard error (n = 8) and as fold induction of control (basal condition, i.e, testes cultured in L-15 only) represented by the dashed line.

**Figure 3.**
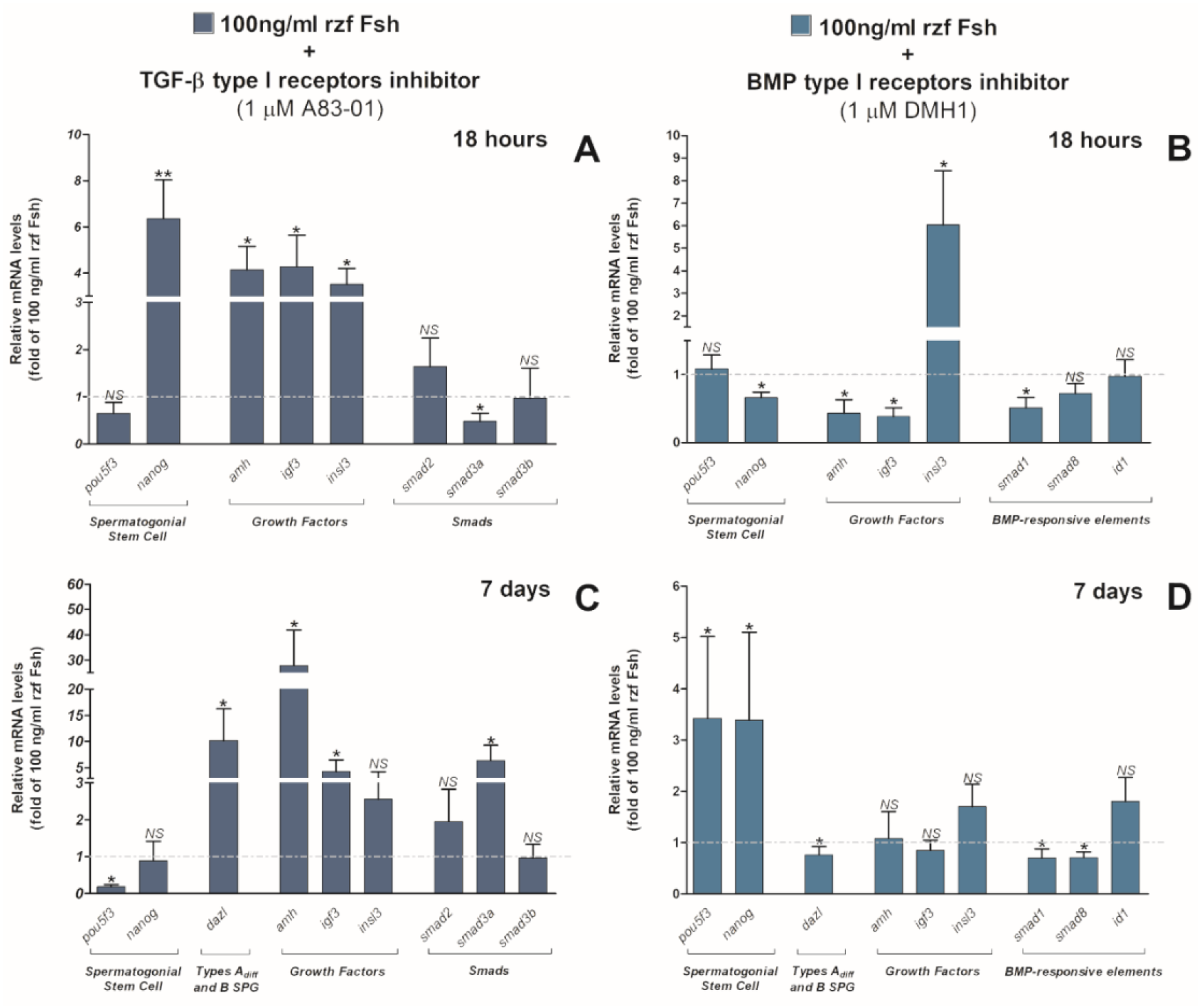
TGF-β and BMP subfamily pathway inhibition effects on gene expression in zebrafish testis tissue cultured under rzf Fsh stimulation. (A-D) Relative expression of spermatogonial stem cell (*pou5f3*, *nanog*), spermatogonial differentiation (*dazl*), growth factors (*amh*, *igf3*, *insl3*) markers, and intracellular messengers of the TGF-β superfamily pathway *(smad1, smad2, smad3a, smad3b, smad8, id1)* in testis tissue cultures co-treated with rzf Fsh (100 ng/mL) and A83-01 inhibitor (1 μM) for 18h (A) and 7 days (C). Co-treatment with rzf Fsh (100 ng/mL) and DMH1 inhibitor (1 μM) for 18h (B) and 7 d (D). Ct values were normalized to the reference gene (β-actin) and expressed relative to the basal group (rzf Fsh). (*) and (**) represent significant differences (p <0,05), (p<0,01), respectively. (NS), no significant differences. Data are presented as the mean ± standard error (n = 8) and as fold induction of control (basal condition, i.e., testes cultivated in rzf Fsh only), represented by the dashed line.

**Figure 4.**
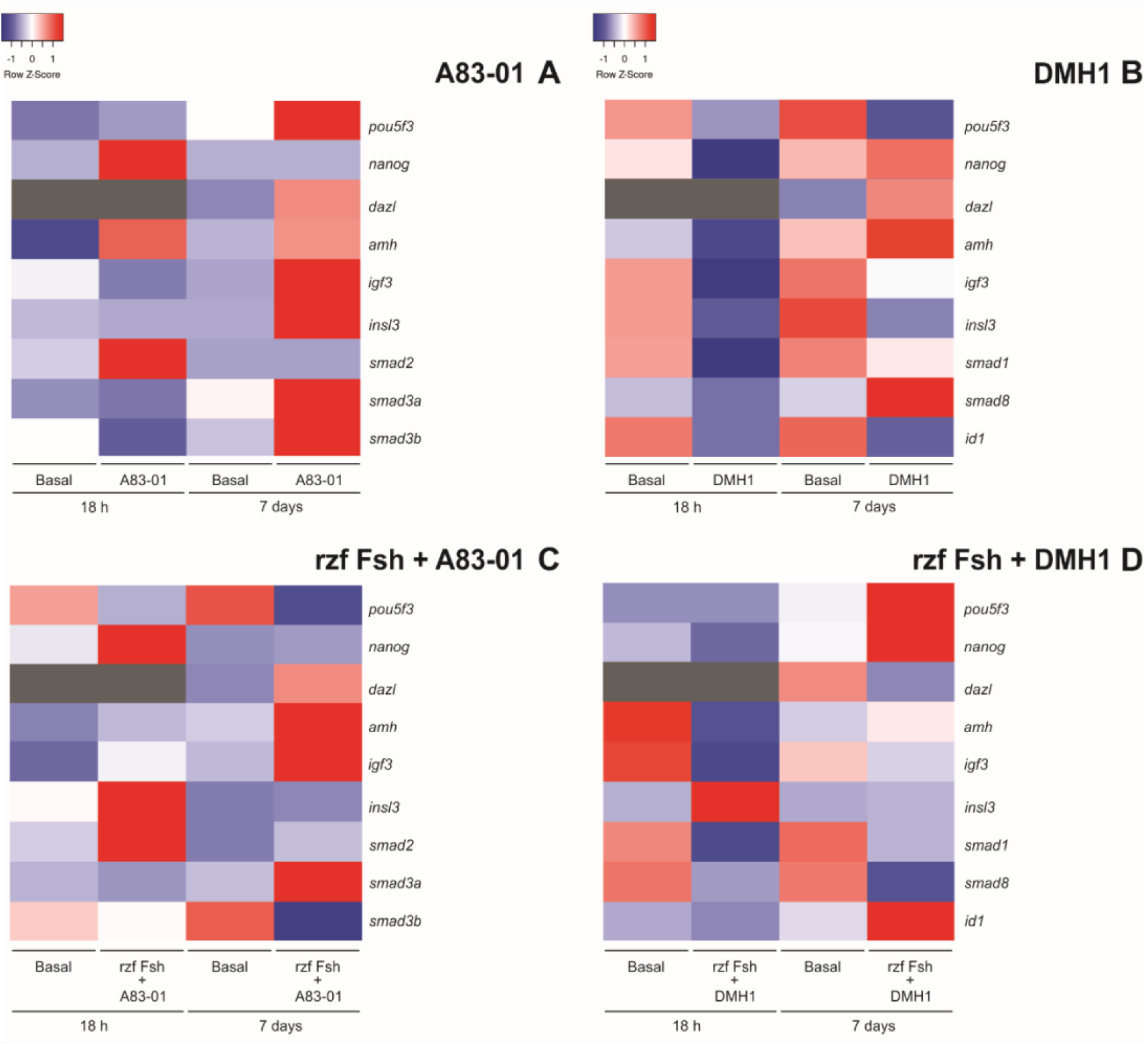
Heat map illustrating TGF-β and BMP subfamily pathway inhibition effects on gene expression in zebrafish testis tissue cultured under basal conditions or rzf Fsh stimulation. (A-D) Heat map illustrating the relative mRNA level of spermatogonial stem cell *(pou5f3, nanog),* spermatogonial differentiation *(dazl),* growth factors *(amh, igf3, insl3)* markers, and intracellular messengers of the TGF-β superfamily pathway *(smad1, smad2, smad3a, smad3b, smad8, id1)* in 18 h and 7 days. Testis tissue cultures treated with the A83-01 inhibitor (1 μM) (A). DMH1 inhibitor (1 μM) (B). Co-treated with rzf Fsh (100 ng/mL) and A83-01 inhibitor (1 μM) (C). Co-treatment with rzf Fsh (100 ng/mL) and DMH1 inhibitor (1 μM) (D). Data shown are log2 values (relative quantification) relative to the average expression. Each colored cell in the heat map represents the standardized relative gene expression value for each sample. The higher expression values are displayed in red, moderate expression values in shades of white (light blue and light red) and lower expression values in blue. Gene expression not evaluated in 18 h (*dazl*) is represented in gray.

In contrast to the 18 h culture, the relative mRNA levels of *igf3, insl3, smad3a* and *smad3b* increased in the testis tissue cultured with A83-01 for 7 days. The expression of the spermatogonial differentiation gene *dazl*, increased significantly following treatment of testis cultures withA83-01 or DMH1 for 7 days (Figures 2, 4). In contrast to 7 days incubation with A83-01, the DMH1 inhibitor reduced *igf3* and *insl3* expression following treatment for the same time period. Overall, compared with 18 h exposure to DMH1, 7 days treatment with DMH1 showed less downregulation activity for all genes analyzed, and a nonsignificant tendency of increase in the relative mRNA levels of *nanog, amh*, and *smad8. id1* was strongly downregulated after 18 h and 7 d of exposure to DMH (Figures 2, 4).

### 3.3 Effects of the A83-01 and DMH1 inhibitors on gene expression of testis tissue cultured in the presence of rzf Fsh

Studies in zebrafish have shown that Fsh acts as an important regulator of the initial phases of spermatogenesis. In particular, stimulation of spermatogonial proliferation and differentiation occurs through the modulation of stimulatory growth factors (e.g., Igf3, Insl3, androgens) and inhibitory growth factors (e.g., Amh). Zebrafish testes were incubated with rzf Fsh (100 ng/mL) in the presence or absence of A83-01 or DMH1 for 18 h and 7 days. The purpose of these experiments was to evaluate the effects of pharmacological inhibition of the TGF-β or BMP subfamily pathways on the testes upon rzf Fsh stimulation. Compared to A83-01 treatment, testis tissue cultured with rzf Fsh and A83-01 for 18 h showed increased primary transcript levels of *nanog, amh, igf3,* and *insl3*, and only *smad3a* was downregulated (Figures 3, 4). After 7 days of culture, co-treatment with rzf Fsh and A83-01 reduced the expression of the spermatogonial stem cell marker *pou5f3*, while up-regulating the spermatogonial differentiation marker *dazl*. (Figures 3, 4). Interestingly, the primary transcripts of the *amh* and *igf3* growth factors continued to be significantly elevated after co-treatment after 7 days (Figures 3, 4). In contrast to co-treatment with rzf Fsh and A83-01 for 18 h, *smad3a* expression increased after co-treatment for 7 days, when compared with its expression in the testes cultured for 7 days with rzf Fsh only (Figures 3, 4). The primary transcript levels of *nanog* and *smad2* did not change in relation to the control group (Figure 3). Similar to 18 h of cultivation with DMH1, 18 h of co-treatment with rzf Fsh and DMH1 predominantly decreased the expression of almost all analyzed genes, with the exception of *insl3* (significantly increased), *pou5f3* (slightly increased, not significant), and *id1*. As observed between DMH1 inhibition for 18 h and 7 days, the 7 days co-treatment with rzf Fsh and DMH1 reduced the downregulation activity compared to 18 h co-treatment (Figures 3, 4). The relative mRNA levels of spermatogonial stem cell markers *(pou5f3* and *nanog)* increased in response to co-treatment with rzf Fsh and DMH1. Differently from 7 days treatment with DMH1 alone, *dazl* was downregulated by co-treatment. Interestingly, *id1* showed unchanged expression at both 18 h and 7 days of co-treatment (Figure 3).

### 3.4 Effects of the A83-01 and DMH1 inhibitors on the zebrafish spermatogonial phase

We evaluated the effects of pharmacological inhibitors of the TGF-β and BMP subfamily pathways on different spermatogonia generations through the frequency of spermatogonial cysts (Aund, Adiff, and B) in testis tissue cultured for seven days (Figure 5). A83-01 treatment increased type B spermatogonia compared to testes cultured under basal condition (L-15 only) (Figure 5A). On the other hand, DMH1 treatment reduced type B spermatogonia frequency and increased the percentage of type A undifferentiated spermatogonia (Aund) compared to the basal condition (L-15 only) (Figure 5B), while undifferentiated spermatogonia showed a slight tendency to decrease with A83-01 treatment (not significantly different) (Figure 5A). Co-treatment with rzf Fsh and A83-01 or DMH1 resulted in opposite cyst frequencies for all early spermatogonial cysts. Co-treatment with rzf Fsh and A83-01 increased the percentage of differentiated spermatogonia and reduced undifferentiated spermatogonia as compared with the contralateral testis exposed with rzf Fsh (Figure 5C, E-H). In contrast, co-treatment with rzf Fsh and DMH1 decreased the frequency of differentiated spermatogonia and augmented undifferentiated spermatogonia compared with testes cultured with rzf Fsh (Figure 5D). The frequency of type B spermatogonia cysts did not change between the groups co-treated with rzf Fsh (Figure 5C and D).

**Figure 5.**
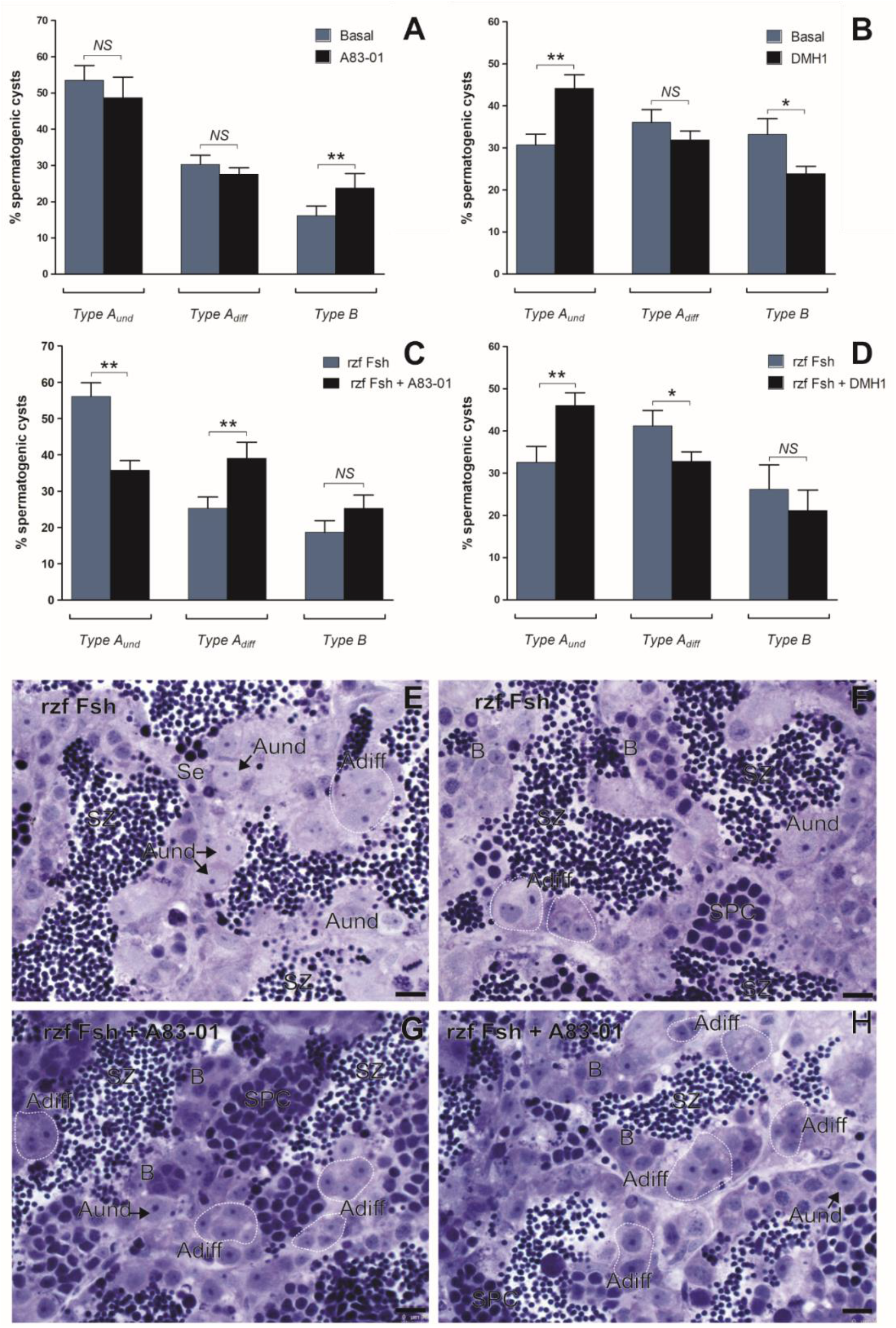
Morphometrical analyses of cultured zebrafish testis tissue after inhibition of TGF-β and BMP subfamily pathways under rzf Fsh stimulation. (A-D) Spermatogenic cyst frequency in zebrafish testis tissue cultured for 7 days. (A and B) Testis tissue cultured under basal conditions (L-15 only) compared to testis tissue cultured in the presence of A83-01 inhibitor (1 μM) (A) or DMH1 inhibitor (1 μM) (B). (C and D) Testis tissue cultured in the presence of 100 ng/mL of rzf Fsh compared to testis tissue co-treated with rzf Fsh (100 ng/mL) and A83-01 inhibitor (1 μM) (C) or rzf Fsh (100 ng/mL) and DMH1 inhibitor (1 μM) (D). (*) and (**) represent significant differences (p < 0,05) and (p < 0,01), respectively, observed between groups for each one of the different types of cysts investigated; (NS) no significant difference. Data are expressed as mean ± standard error (n = 6). (E-H) Morphology of the zebrafish testis tissue cultured for 7 days. (E and F) testis incubated in the presence of rzf Fsh (100 ng/mL) or (G and H) with the presence of rzf Fsh (100 ng/mL) and 1 μM of A83-01. Blue toluidine staining. The white dotted line delimits the differentiation Type A spermatogonia cysts (Adiff). Aund = undifferentiated type A spermatogonia cysts; B= type B spermatogonia cysts; SPC= spermatocyte cysts; SZ= spermatozoa; SE=Sertoli cells. Scale bar=10 μm.

### 3.5 Basal and Fsh-induced expression of smads and id1 in testis tissue culture

We examined *smad* expression in testis tissue cultured under basal conditions (L-15). Interestingly, *smad2* showed higher expression in relation to the others *smads* of the TGF-β subfamily pathway (Figure 6A). The *smads* of the BMP subfamily pathway *smad1* and *smad8* were unchanged, however, they were expressed at higher levels compared to *smad3b* (Figure 6A). *smad3b* was less expressed in comparison to all *smads* analyzed, with no significant difference compared to its isoform *smad3a* (Figure 6A). We also evaluated the effects of rzf Fsh on the expression of *smads* and *id1* in cultured zebrafish testis tissues (Figure 6B). The rzf Fsh was only able to increase the expression of *smad3b*, no other Smad analyzed responded to rzf Fsh stimulation. The *id1* gene was downregulated by rzf Fsh stimulation (Figure 6B).

**Figure 6.**
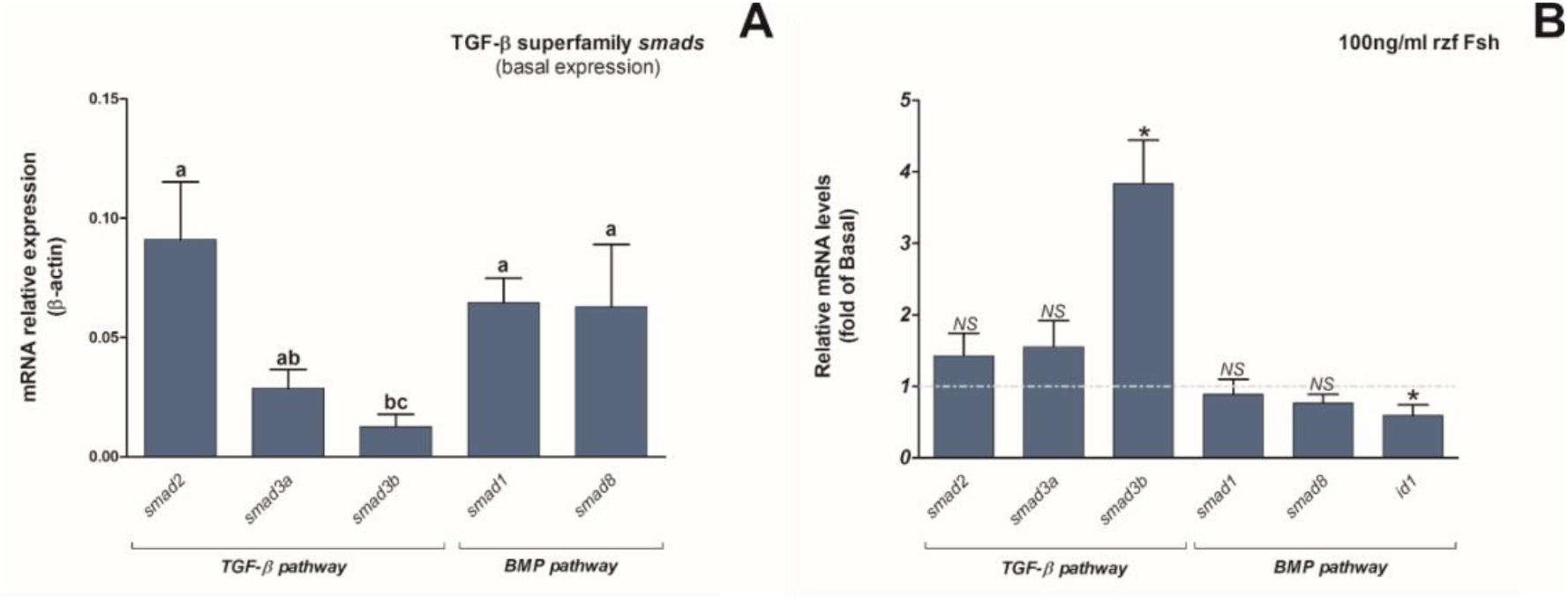
Expression of *smads* under basal conditions (L-15 only) and following stimulation with rzf Fsh in cultured zebrafish testis tissue. Relative expression of the TGF-β-superfamily *smads* in zebrafish testis tissue cultured with L-15 (A) or rzf Fsh (100 ng/ml) (B) for 18 h. (A) The Cts were normalized by the reference gene (β-actin) and expressed relative to the basal group (L-15 only) or (B) relative to the dCTs mean of all the *smads* evaluated. (*) significant difference (p < 0,05); (NS). no significant difference. (A) Data are presented as the mean ± standard error (n = 8) and values labeled with different letters are significantly different (p < 0,05) (B) Data are presented as the mean ± standard error (n = 8) and as fold induction of control (basal condition, i.e, testes tissue cultured in L-15 only) represented by the dashed line.

## 4. Discussion

Pituitary gonadotropins (Fsh and Lh) act as endocrine regulators of testicular function, stimulating testicular development through production of growth factors and sexual steroids [43]. Although Fsh acts mainly in the initial stages of spermatogenesis, Lh plays a role in the final stages [43]. Recent studies on fish spermatogenesis demonstrated that Fsh promotes the proliferation and differentiation of spermatogonial cells by increasing the production of stimulatory growth factors such as Igf3 [32,37,44–46]. Furthermore, Fsh decreased the expression of *amh*, a member of the TGF-β superfamily that inhibits spermatogonial differentiation [47]. Moreover, recent RNAseq studies have shown that Fsh also regulates *insl3* in Leydig cells and several TGF-β superfamily genes in zebrafish testis tissue culture, such as *inha, inhbab, lft1, bmp7b, nog2, and gsdf* [45].

Considering the role of the Fsh-mediated TGF-β superfamily in zebrafish spermatogenesis, we evaluated the role of TGF-β and BMP subfamily pathways in zebrafish spermatogenesis using the pharmacological inhibitors A83-01 and DMH1. Tojo and Collaborators [26] synthesized and characterized A83-01 to block as many TGF-β subfamily pathway receptors as possible. They found efficiency in inhibiting TGF-β type I receptors (Alks 4, 5, and 7) at a concentration of 1 μM in mink lung epithelial (Mv1Lu) cell culture; a concentration ten times lower than that required for the previous inhibitor (SB-431542) [48] to have a similar effect [26]. DMH1 is a specific BMP Type I receptor inhibitor with no off-target effects [27]. The previously used BMP Type I receptor inhibitor compounds Dorsomorphin (Compound C) and LDN-193189 showed off-targets effects inhibiting some TGF-β Type I receptors and many others unspecific kinases [21,27,49]. Thus, DMH1 was more effective in inhibiting the BMP subfamily. A previous experiment using DMH1 examined the concentration required to inhibit the BMP subfamily pathway [29]. However, only 3 μM and 5 μM were used to verify the *Id1* gene response in NSCLC cell culture. For this reason, we performed a concentration test based on *id1* gene downregulation and found BMP type I receptor inhibition with only 1 μM DMH1. Similarly, *id1* expression was successfully suppressed following treatment of adult zebrafish during fin regeneration [50] and NSCLC cells [29] with 10 and 3 μM, respectively. In addition, inactivating mutations in ALK6 (BMP type I receptor) in zebrafish adult testes led to *id1* downregulation [51].

In zebrafish embryos, TGF-β subfamily smads (smad2, smad3a and smad3b) are mainly maternally inherited and have different patterns of expression in the blastula and gastrula stages [52,53]. Similarly, BMP subfamily *smads (smad1* and *smad5*) are differentially expressed in zebrafish embryos, smad5 is maternally inherited, and *smad1* mRNA is undetectable prior to mid gastrulation at cleave, blastula, or early gastrula stages [54,55]. However, there is less information regarding *smad8,* and we did not find any studies on its expression pattern during zebrafish embryonic development. In mouse, *smad8* expression is initially confined to the visceral endoderm and shows a tightly regulated pattern of expression restricted to only a few tissue sites [56]. In contrast, *smad1* and *smad5* are ubiquitously expressed throughout the mouse embryo [56].

Despite the studies performed with zebrafish embryos, as far as we know, *smads* expression in zebrafish testes was not previously evaluated. Here, in adult zebrafish testis tissue cultured in basal condition (L-15 only), the levels of the primary transcripts of *smad2* were higher than those of *smad3b*. Similar results were found comparing *Smad2* to *Smad3* in the developing and adult rat testes [22]. We did not observe any discrepancy in expression among *the smads*. The same balanced pattern of expression was reported for the TGF-β and BMP subfamily *Smads* in rat and mouse testes [1,22]. Western blot and fluorescent immunohistochemistry showed that Smad2 and Smad3 had parallel expression in the postnatal stages of developing rat testes and had cell type-specific expression in adult testes. Although *Smad2* and *Smad3* are differentially expressed according to the spermatogenesis cell type in adult rats, both *Smad2* and *Smad3* are expressed in all seminiferous tubules [22]. In the mouse testis, *in situ* hybridization analyses for the BMP subfamily *Smads* showed that *Smad1* is more expressed in initial spermatogenic cells (spermatogonial stem cells and differentiated spermatogonia) and *Smad8* shows higher expression in late spermatogenesis (spermatocyte and spermatid) presenting an alternating and balanced pattern of expression throughout the spermatogenesis process [1].

Following cultivation with rzf Fsh (100 ng/mL) for 18 h, only the levels of *smad3b* were increased among all *smads* examined. In contrast, *id1* expression was downregulated by rzf Fsh. These results showed that direct or indirect regulation of Fsh can change the intracellular levels of TGF-β superfamily members. This also suggests that these pathways are recruited by rzf Fsh downstream signaling, corroborating the RNAseq data published previously [45]. The use of pharmacological inhibitors of TGF-β superfamily members in zebrafish testis tissue cultured in the absence or presence of rzf Fsh for 18 h or 7 days affected the expression of *smads* differently. *smad2* mRNA levels remained unaltered by the A83-01 inhibitor under all conditions. Expression of *smad3a* and *smad3b* showed higher variation among the groups in a very interesting way. The inhibition with A83-01 for short-term (18 h) only inhibited *smad3b* transcription (significantly) and *smad3a* (not significantly), and long-term (7 days) treatment increased expression of these two *smads* (*smad3a* e *smad3b*). Similarly, following co-treatment with rzf Fsh and A83-01, *smad3a* transcripts decreased in the short term (18 h) and increased after prolonged incubation (7 days). These results show that in the short term, the inhibitor A83-01 suppress *smad3a* and *smad3b* expression (depending on the condition); however, in the long term, even with pharmacological block, the cells produced more transcripts of these molecules. This cellular response can be a compensatory mechanism to maintain homeostasis in the TGF-β subfamily pathway.

The inhibition of the BMP subfamily pathway resulted in a strong reduction in BMP-responsive component (*smad1, smad8*, and *id1*) expression at 18 h and 7 days, with the exception of *smad8* at 7 days. Despite *smad1* reduction at 7 days, its reduction was less pronounced when compared with that at 18 h, furthermore *smad8* showed increased expression (not significant) after 7 days. This could indicate the same compensatory activity reported in the inhibition of TGF-β subfamily *smads*. Unlike treatment with only DMH1, co-treatment with rzf Fsh and DMH, had no effect on the levels of *id1* at 18 h and 7 days. This unexpected result could be explained by the fact that *id1* showed reduced expression following treatment with rzf Fsh when compared with the basal condition (l-15 only), and the reduction already caused by rzf Fsh in control groups can mask the downregulation effect of the DMH1 inhibitor.

We did not evaluate the phosphorylation of Smads, which indicates their activation. However, the decreased expression of *smad3a*, *smad3b* and *smad8* at short-term and *smad1* at short-term and long-term supports the efficiency of the inhibitor in repressing the pathway.

We also analyzed the effect of inhibitors on the expression of relevant genes for spermatogenesis, such as spermatogonial stem cell (*pou5f3* and *nanog*), spermatogonial differentiation (*dazl*), and growth factor (*amh*, *igf3*, and *insl3*) genes. We did not measure *dazl* expression in the short term, since 18 h could be insufficient to evaluate a differentiation marker. Leal et al. [35] concluded that, in cystic spermatogenesis, it is possible to determine time elapsed between preleptotene/leptotene and spermatozoa. However, the duration of spermatogonial differentiation cannot be deduced.

The short-term inhibition of the BMP subfamily pathway showed a massive suppression effect on the expression of spermatogonial stem cell, growth factor, and BMP-responsive element genes. The simultaneous suppression of spermatogonial stem cell and pro-differentiation growth factors (*igf3* and *insl3*) genes showed that in the short term, the inhibition of the BMP subfamily pathway suppresses initial steps of spermatogonial phase (late spermatogenesis stages were not evaluated), reducing both spermatogonial self-renewal and differentiation. Compared to short-term inhibition of the TGF-β subfamily pathway, BMP type I receptor inhibition had a greater impact on the expression of testis genes. The clear reduction of early spermatogenesis markers and stimulatory growth factors suggests that the BMP subfamily pathway is more active in the early spermatogenesis stages and may be more fundamental to testis function in this stage than the TGF-β subfamily pathway. Morphological analysis was not performed following treatment with A83-01 and DMH1 for 18 h because few effects would be seen in this short time since the complete process of spermatogenesis in zebrafish takes approximately 6 days [35].

Prolonged inhibition of the TGF-β subfamily pathway with A83-01 stimulates the expression of *igf3* and *insl3*, important growth factors of spermatogonial pro-differentiation, and *dazl,* a gene marker of type Adiff and B spermatogonia [32]. Igf3, as anteriorly mentioned, stimulates spermatogonial differentiation in the zebrafish testes [32]. Insl3 (insulin-like peptide 3) is produced by Leydig cells and promotes spermatogonial differentiation in zebrafish [57]. Furthermore, zebrafish testis tissue cultured with rzf Igf3 (100 ng/mL) demonstrated elevated levels of *dazl* [32]. As a consequence of this environment favoring differentiation, more differentiated cells are expected to be found in the spermatogenesis process. Histomorphometrical analysis showed that despite type Adiff spermatogonia not being altered by the inhibitor, type B spermatogonia cysts showed a strong increase.

Prolonged inhibition of the BMP subfamily pathway by DMH1 showed some intriguing results. *dazl* was strongly upregulated by the inhibitor despite downregulation of differentiation growth factors (*igf3* and *insl3*). Differently, Nóbrega and collaborators [32] showed, in zebrafish testis tissue cultured for 7 days, that rzf Igf3 (100 ng/mL) increased *dazl*, while a low rzf Igf3 dose (10 ng/mL) caused *dazl* downregulation. This shows that with BMP subfamily pathway inhibition, *dazl* stimulation is not mediated by Igf3 growth factor. Histomorphometrical analysis in the testes with BMP subfamily inhibited showed an increase in type A undifferentiated spermatogonia, similar to *Dorsomorphin* in zebrafish testis tissue culture [58]. However, Wong and Collodi [58] did not evaluate the type A differentiated spermatogonia in zebrafish testis.

We have some unclear histomorphometrical results regarding the effects of treatment with A83-01 or DMH1 for 7 days. Despite the fact that type B spermatogonia responded to an increase in differentiation growth factors mediated by A83-01 compared with the basal group, type Adiff spermatogonia did not respond to differentiation stimuli. In addition, in long-term exposure, DMH1-mediated downregulation of *pou5f3* cannot be explained because frequency of type A undifferentiated spermatogonia are increased according to histomorphometric analyses, and upregulation of *dazl* was followed by a decrease in differentiated spermatogonia (Adiff, not significant; and type B, significant); however, these reductions in differentiated spermatogonia are expected with downregulation of differentiation growth factors. Therefore, this unusual relationship between expression and morphology might occur because of the lack of regulatory factors, which may be insufficient for *in vitro* experiments. These regulatory factors could be dependent on endocrine stimulation by Fsh and Lh, the major regulatory hormones of spermatogenesis. Corroborating this hypothesis, in the presence of rzf Fsh no discrepancies between the expression and morphology were observed, as shown below.

Considering that rzf Fsh stimulates spermatogonial proliferation and creates a pro-differentiation environment by increasing Igf3, Insl3, and androgens [32,45,57], it is hypothesized that co-treatment with rzf Fsh and A83-01 or DMH1 inhibitors will have an accentuating and additive effect in creating a pro-differentiation environment. The gene expression analysis of the testis tissue cultured for 18 h or 7 days with rzf Fsh in the presence of the A83-01 inhibitor corroborated this hypothesis. Inhibition of the TGF-β subfamily pathway with rzf Fsh for 18 h enhanced the pro-differentiation (*igf3* and *insl3*) and pro-proliferation (*nanog* and *amh*) environment. Otherwise, Morais and collaborators [46] showed that treatment with rzf Amh (10 μg/mL) for 3 days reduced *igf3* and *insl3* expression in zebrafish testis tissue culture. Also, Skaar and collaborators [47] found that recombinant Amh downregulated the stimulatory effect of Fsh on *insl3*. In contrast, following TGF-β subfamily pathway inhibition with rzf Fsh for 7 days, *amh* increase was accompanied by *igf3* and *insl3* (not significant) upregulation. This indicates that the TGF-β subfamily pathway might be indirectly involved in Amh-mediated downregulation of *igf3* and *insl3* in zebrafish testes.

Following co-treatment of TGF-β subfamily pathway inhibition with rzf Fsh for 7 days, elevated levels of *igf3* and *dazl* expression were observed. Interestingly, the expression of the spermatogonial stem cell marker *pou5f3* decreased. This suggests a reduction in stem cell function due to exacerbated spermatogonial differentiation. The histomorphometrical analysis of the testis tissue cultured with A83-01 and rzf Fsh for 7 days were in accordance with the gene expression data, showing an increase in the differentiated spermatogonia (Adiff) and reduction of undifferentiated spermatogonia (Aund). Thus, the TGF-β subfamily pathway plays a role in the self-renewal and proliferation of spermatogonial stem cells in zebrafish. These findings are consistent with previous studies that utilized the SB431542 inhibitor, an inhibitor of the TGF-β subfamily pathway in testis tissue cultures during mouse fetal development [59,60]. In both studies, the inhibition caused an increase in the expression of meiosis marker genes *(Stra8, Rec8*, and *Dmc1*[60]) and *(Stra8, Rec8, Dmc1* and *Mszx1*[59]). Souquet et al. [59] found that in XY gonads cultivated with SB431542 inhibitor, typical morphological features of meiosis were observed in germ cells compared with the basal group. In the same study, recombinant Nodal significantly reduced *Stra8* expression and germ cell meiosis in cultured 11.5 days post-conception ovaries, reinforcing the conclusion that the TGF-β subfamily pathway prevents meiosis in embryonic germ cells. Similar to the present study, these results suggest that the TGF-β subfamily pathway plays a role in spermatogonial proliferation to the detriment of differentiation.

BMP subfamily pathway inhibition with DMH1 (only) for 18 h and 7 days showed the majority of gene repression. The same was observed for BMP subfamily pathway inhibition with DMH1 and rzf Fsh for 18h. BMP subfamily inhibition for 18 h, independent of Fsh presence or absence, seems to be more aggressive to zebrafish testis than TGF-β subfamily pathway inhibition with A83-01; both proliferation and differentiation were impaired.

In contrast to the expected effects of rzf Fsh, long-term co-treatment with DMH1 and rzf Fsh did not show pro-differentiation activity. The long-term inhibition of the BMP subfamily pathway with rzf Fsh resulted in an increase in spermatogonial stem cell markers and downregulation of *dazl*, indicating stimulation of undifferentiated spermatogonia proliferation and decreased differentiation. These effects on early spermatogenesis were also observed in histomorphometric analysis; increased the percentage of type Aund, and decreased type Adiff. Inhibition of BMP subfamily pathway (DMH1 only) strongly downregulates differentiation growth factors *(igf3* and *insl3*). This inhibitory effect was counterbalanced by rzf Fsh, which minimized inhibition of the differentiation growth factors promoted by DMH1 but still decreased spermatogonial differentiation.

Thus, the BMP subfamily pathway in zebrafish mainly promotes spermatogonial differentiation. Wong and Collodi [58], using *dorsomorphin,* BMP subfamily pathway inhibitor, observed induction of spermatogonial proliferation in zebrafish testes. The same effect was reported in zebrafish spermatogenesis by Neumann et al. [51] through the mutation of the ALK6 receptor of the BMP subfamily pathway. These morphological results can also be explained by the BMP subfamily pathway, which inhibits spermatogonial proliferation. However, following DMH1 treatment alone (18 h and 7 days), all undifferentiated spermatogonial markers were strongly downregulated, with the exception of *nanog* after 7 days. Thus, blocking the two branches of the TGF-β superfamily pathway points to an antagonistic role between the TGF-β and BMP subfamily pathways.

To observe the antagonistic effect between TGF-β and BMP subfamily pathways, Fsh is a useful stimulus because it can accentuate the functions of the pathway that is not blocked, making the differences between them more evident. In zebrafish testes, Fsh inhibits the expression of *ift1* (Lefty1, antagonist of Nodal) and *nog2* (Noggin 2, antagonist of BMPs) [45], increasing the availability of ligands for their respective type I receptors. Fsh regulates the two branches of TGF-β superfamily pathways differently. Among the TGF-β subfamily members, Fsh stimulation repressed *inhbab* (activin A) expression in zebrafish [45] and rainbow trout testes *in vitro* [61] and upregulated *inha* (inhibin) in zebrafish and rainbow trout testes *in vitro* [45,62]. Among the BMP subfamily members, Fsh downregulates *amh* in zebrafish testis tissue cultures [32]. Few studies have evaluated Fsh effects on Bmp expression in testes, and Crespo and collaborators [45] found upregulation of *bmp7b* in zebrafish testes following treatment with Fsh. However, this effect was only observed after re-starting spermatogenesis, which was interrupted by estradiol. Estradiol promotes androgen insufficiency and type A undifferentiated spermatogonia enrichment. We did not find data concerning Fsh regulation of TGF-β (1-3) or TGF-β functions in adult testes, especially in fish. TGF-β (1-3) is related to PGCs (primordial germ cells) migration in the developing testes [1,8]. However, studies on the adult gonads that determine their functions are lacking.

Following TGF-β subfamily pathway inhibition by rzf Fsh, Bmps may mediate differentiation through the BMP subfamily pathway *in vitro*. rzf Fsh upregulated *amh*, an inhibitor of differentiation. Screening of BMPs in all mouse testis stages found that *Bmp2, Bmp3, Bmp3b, Bmp4, Bmp5, Bmp6, Bmp7, Bmp8a, Bmp8b* and *Bmp15* are expressed [63]. However, studies to elucidate their functions in adult testes are still lacking. The functions of BMP4, BMP7, BMP8a, and BMP8b are better understood in adult testis spermatogenesis; BMP4 acts as a differentiation factor in initial mouse testis spermatogenesis [1,8,64] whereas BMP7, BMP8a, and BMP8b act in meiotic cells [65]. Therefore, BMP4 is a candidate mediator of differentiation through the BMP subfamily pathway during early spermatogenesis, promoting the transition of undifferentiated spermatogonia to differentiated spermatogonia during TGF-β subfamily pathway inhibition.

Following BMP subfamily pathway inhibition with rzf Fsh, Nodal is the probable mediator of spermatogonial self-renewal through the TGF-β subfamily pathway *in vitro*.Inhibin regulates endocrine function in the pituitary [8,66] and Activin is not related to the stimulation of self-renewal in adult testes. Activin A is considered an inhibitor of differentiation; in postnatal mouse testis the transcripts of the meiotic germ cell markers (*Sycp3* and *Ccnd3*) were upregulated by *Inhba* knockin (replacing original sequences for *Inhbb,* a less bioactive Activin) and morphological analyzes found increased number of spermatocytes in relation to spermatogonia. Activin A seems to block differentiation in the meiotic stage because *Inhba* knockin enhanced capacity of type B spermatogonia to transition into spermatocytes [67].

In mouse postnatal testes *in vitro*, Activin A acts in early spermatogenesis, reducing differentiated spermatogonia and repressing the pro-differentiation effects of FSH [68]. During BMP subfamily pathway inhibition, Activin A may mediate the reduction in differentiated type A spermatogonia induced by the TGF-β subfamily pathway through the repression of the differentiation effects of FSH. Nodal has been described as a promoter of self-renewal in developing mouse testes [69] and several other tissues [70]. Thus, Nodal is a probable mediator of the pro-self-renewal effects of the TGF-β subfamily pathway, increasing undifferentiated spermatogonia during BMP subfamily pathway inhibition. In addition, Fsh facilitates Nodal activity by downregulating its antagonist, *lft 1* [45]. TGF-βs are associated with quiescence in some tissues [3,41] and are produced by the Sertoli cells. Since Fsh stimulates functions in Sertoli cells, it is possible that Fsh can stimulate TGF-β production, thereby stimulating pluripotency through the TGF-β subfamily pathway. Future studies that specifically block candidate ligands concomitantly with inhibitors may contribute to the clarification of these questions.

DMH1 inhibition promotes an intense reduction in spermatogenesis-related gene expression. However, a great recovery in the transcriptional activity of these genes was observed after co-treatment with DMH1 and rzf Fsh. The same effect was not observed with A83-01 treatment compared to co-treatment with A83-01 and rzf Fsh. This evidence suggests that Fsh may have a greater capacity to stimulate TGF-β subfamily members instead of BMP subfamily members in early spermatogenesis.

An interesting result was the increased expression of the *amh* gene in testis tissue cultured with rzf Fsh and A83-01 for 7 days. This observation seems contradictory because previous studies have shown that Fsh decreases *amh* expression [46,47]. The relationship between *amh* and *igf3* expression in these cultures was also unforeseen because both genes in previous studies showed an inversely proportional gene expression [32,46]. Inhibition of the TGF-β subfamily pathway in the presence of rzf Fsh creates an environment of excessive differentiation, leading to a decrease in spermatogonial stem cells, suggesting that the increase in *amh* expression acts as a way to block differentiation and stimulate spermatogonial stem cell proliferation. Supporting this hypothesis, we found no variation in *amh* gene expression in cultures without rzf Fsh stimulation.

Another interesting result concerns *insl3* response to rzf Fsh. *insl3* did not respond to treatment with inhibitors (A83-01 or DMH1) and rzf Fsh for 7 days, in contrast to previous studies where Fsh upregulated *insl3* expression in zebrafish testis after 48 h incubation [44,45]. However, when utilized 18 h of exposure for both inhibitors (A83-01 or DMH1) and rzf Fsh, *insl3* was upregulated. It is possible that in long-term exposure (7 days), which covers complete spermatogenesis in zebrafish (approximately 6 days) [33], *insl3* is not upregulated by Fsh.

## Conclusion

Our results showed that the TGF-β subfamily pathway favors spermatogonial proliferation in adult zebrafish testis (Figure 7). When using a pharmacological inhibitor of this pathway, A83-01, together with Fsh, we demonstrated that it accentuates the pro-differentiation effects of Fsh, excessively stimulating spermatogonial differentiation and reducing the pool of spermatogonial stem cells (Figure 7). An interesting result was the increased expression of *amh* under these conditions, suggesting that it may block excessive differentiation and simultaneously promote spermatogonial stem cell proliferation. The BMP subfamily pathway promotes spermatogonial differentiation in adult zebrafish testes. In co-treatment with Fsh and DMH1, the reduction in differentiated spermatogonia is clear. BMP subfamily inhibition in the short term, independent of the presence or absence of Fsh, appears to be more aggressive in zebrafish testes than TGF-β subfamily inhibition. The BMP subfamily pathway is more stimulatory during initial spermatogenesis, whereas the TGF-β subfamily pathway is more inhibitory. In summary, the TGF-β subfamily pathway promotes spermatogonial proliferation or inhibits spermatogonial stem cell differentiation, whereas the BMP subfamily pathway promotes spermatogonial differentiation (Figure 7).

**Figure 7.**
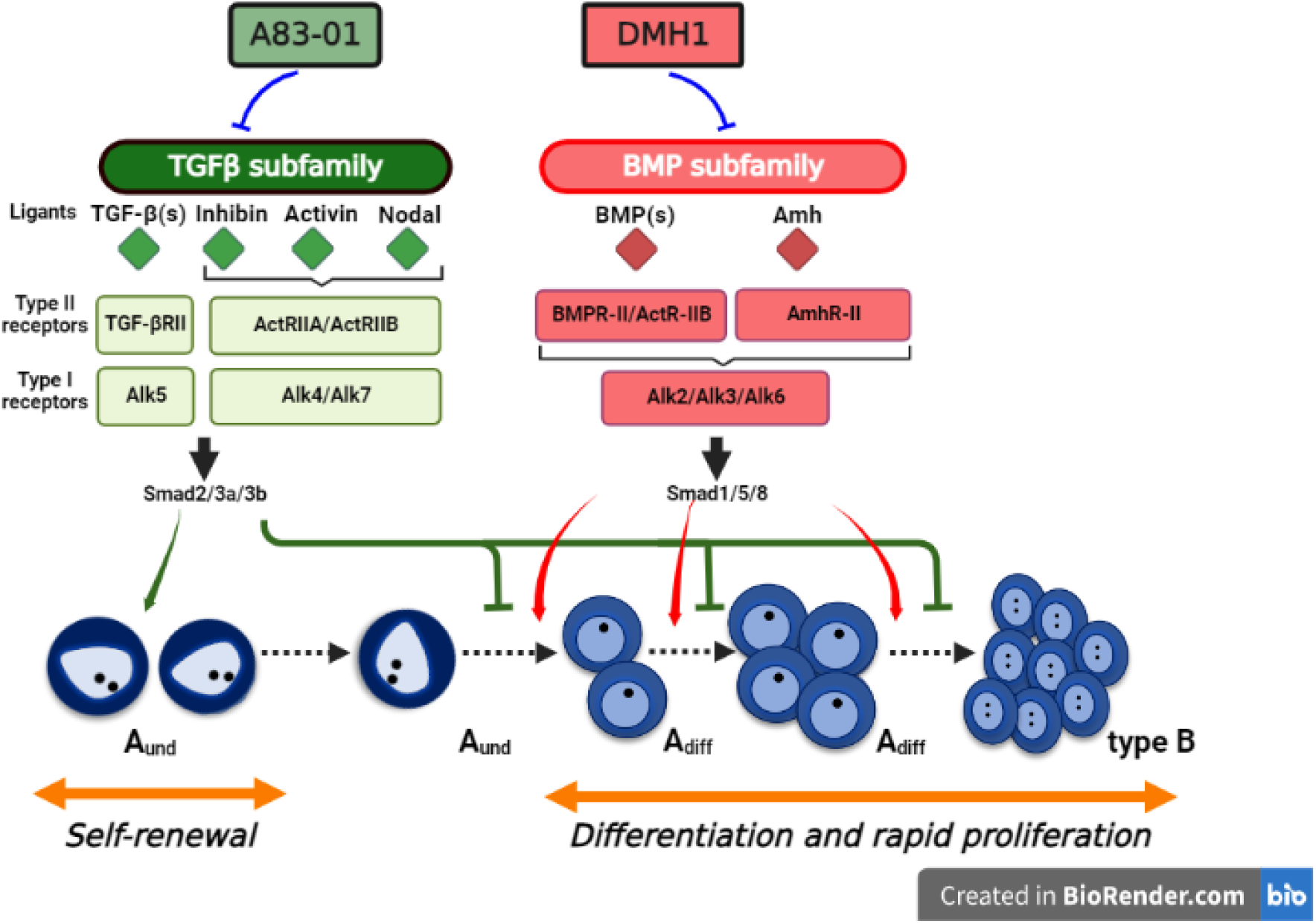
Schematic illustration of TGF-β superfamily signaling pathway in the zebrafish spermatogonial activity (self-renewal *vs*. differentiation). The TGF-β superfamily signaling pathway is subdivided into TGF-β and Bone morphogenetic proteins (BMP) subfamilies. The roles of TGF-β and BMP subfamily pathways were addressed through A83-01 and DMH1 inhibitors, respectively. Our results showed that A83-01 potentiated the follicle-stimulating hormone (Fsh) effects on zebrafish spermatogenesis, reducing type A undifferentiated spermatogonia (Aund) and increasing differentiated spermatogonia (type Adiff and type B spermatogonia). For the BMP signaling pathway, exposure to DMH1 inhibitor showed opposite effects as compared to TGF-β superfamily signaling pathway inhibitor. In conclusion, our study demonstrated that TGF-β and BMP subfamily pathways exert a role in zebrafish spermatogonial niche with antagonistic functions for the spermatogonia fate. The TGF-β subfamily pathway is involved with spermatogonial self-renewal and inhibition of differentiation, whereas the BMP subfamily pathway promotes spermatogonial differentiation and their rapid proliferation.

## Author Contributions

R.H.N and D.F.C., designed the study; D.F.C, J.M.B.R, M.S.R., M.A.O and L.B.D performed the experiments; D.F.C and R.H.N analyzed the data; D.F.C and R.H.N wrote the paper. All authors edited the article.

## Funding

This research was supported by São Paulo Research Foundation (FAPESP) (grant numbers 2017/15793-7 - granted to M.S.R; 2020/03569-8 – granted to R.H.N; 2019/05643-3 granted to L.B.D; 2018/16595-7 – granted to J.M.B.R; 2014/25313-4 granted to M.A.O); and financed in part by the Coordenação de Aperfeiçoamento de Pessoal de Nível Superior—Brasil (CAPES)—Finance Code 001 (granted to D.F.C). R.H.N was awarded productivity scholarship from the Brazilian National Council for Scientific and Technological Development (CNPq) (proc. nos. 305808/2020-6)

## Notes

### Competing Interest Statement

The authors have declared no competing interest.

